# Developmental hourglass and heterochronic shifts in fin and limb development

**DOI:** 10.1101/2020.01.10.901173

**Authors:** Koh Onimaru, Kaori Tatsumi, Chiharu Tanegashima, Mitsutaka Kadota, Osamu Nishimura, Shigehiro Kuraku

## Abstract

How genetic changes are linked to morphological novelties and developmental constraints remains elusive. Here we investigate genetic apparatuses that distinguish fish fins from tetrapod limbs by analyzing transcriptomes and open chromatin regions (OCRs). Specifically, we compare mouse forelimb buds with pectoral fin buds of a slowly evolving species, the brown-banded bamboo shark (*Chiloscyllium punctatum*). A transcriptomic comparison with an accurate orthology map reveals both a mass heterochrony and hourglass-shaped conservation of gene expression between fins and limbs. Furthermore, chromatin-accessibility data indicate that conserved regulatory sequences are most active during mid-stage limb development. During this stage, stage-specific and tissue-specific OCRs are also enriched. Together, early and late stages of fin/limb development are more permissive to mutations, which may have contributed to the major morphological changes during the fin-to-limb evolution. We also hypothesize that the middle stages are constrained by regulatory complexity that results from dynamic and tissue-specific transcriptional controls.

## Introduction

The genetic control of morphological diversity among multicellular organisms is a central interest in evolutionary biology. In particular, our understanding of how morphological novelties are linked to the emergence of their respective genetic apparatuses is limited^1^. In addition, it is still unclear to what extent internal constraints, such as pleiotropy, affect evolvability^2^. The fin-to-limb transition is a classic, yet still influential, case study that contributes to our understanding of morphological evolution. In general, tetrapod limbs are composed of three modules, the stylopod, zeugopod, and autopod, which are ordered proximally to distally (Fig. 1a). In contrast, fish fins are often subdivided into quite different anatomical modules along the anterior–posterior axis—the propterygium, mesopterygium, and metapterygium (Fig. 1a). Although it is still controversial how this different skeletal arrangement compares with the archetypal tetrapod limb, the autopod (wrist and digits) seems to be the most apparent morphological novelty during the fin-to-limb transition ^3^. Despite intensive comparative studies of developmental gene regulation, genetic machinery that differs between fins and limbs remains elusive. Instead, several studies revealed that autopod-specific regulation of *Hoxa13* and *Hoxd10–13*, which control autopod formation, is also conserved in non-tetrapod vertebrates^4–6^. Whereas some tetrapod-specific gene regulation has been proposed, these studies were narrowly focused on *Hox* genes^7–10^. Therefore, a genome-wide systematic study is required to identify the genetic differences between fish fins and tetrapod limbs.

**Fig. 1.**
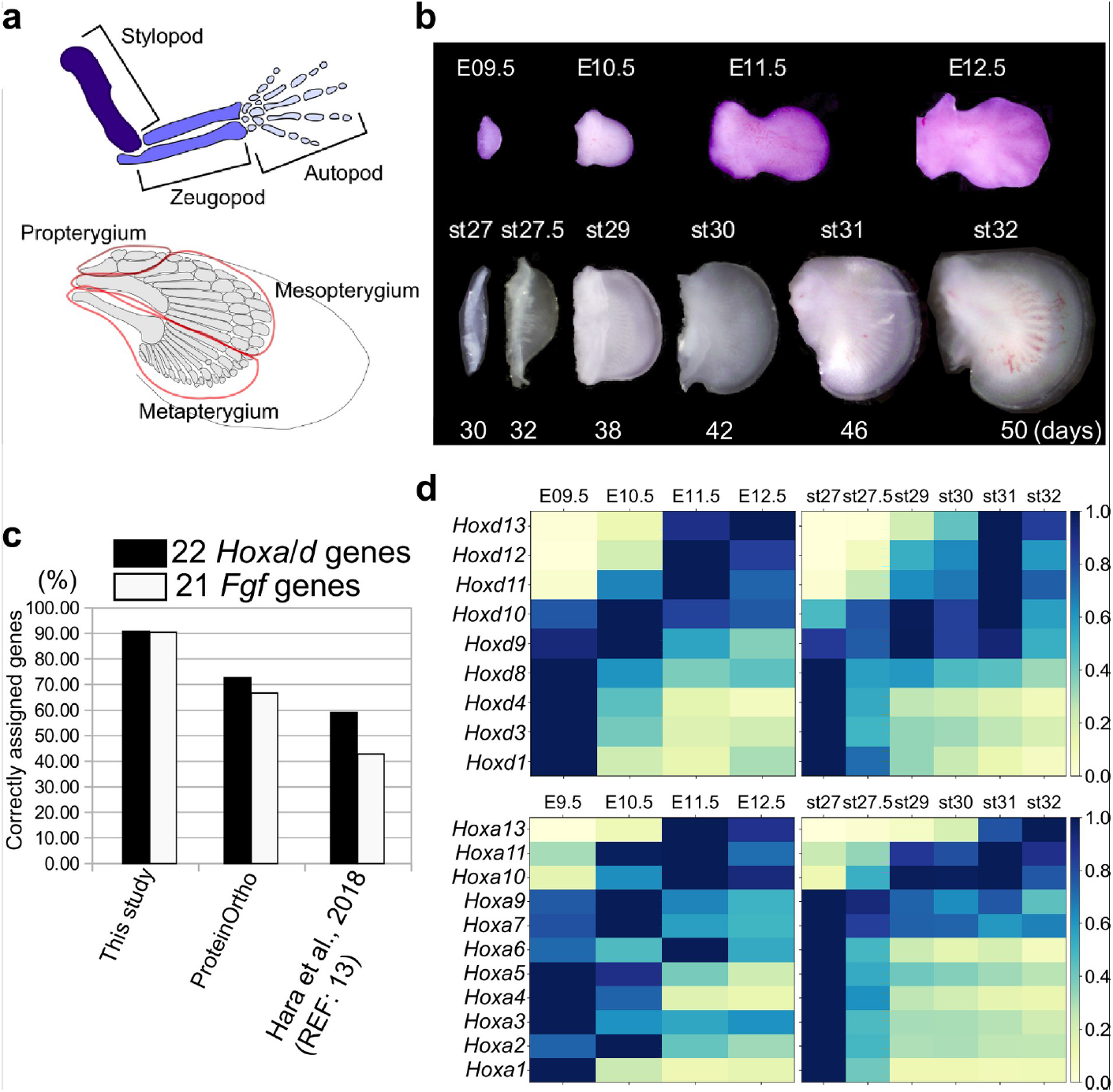
Transcriptome analysis and orthology assignment. **a**, The skeletal patterns of a mouse limb (top) and a bamboo shark pectoral fin (bottom). Anterior is to the top; distal is to the right. **b**, Mouse forelimb buds and bamboo shark pectoral fin buds that were analyzed by RNA-seq. **c**, Comparison of the accuracy of three orthology assignment methods. Vertical axis, the percentages of correctly assigned *Hoxa* and *Hoxd* paralogs (black bars) and *Fgf* paralogs (white bars). **d**, Heat map visualization of the transcription profile of *Hoxa* and *Hoxd* genes in mouse limb buds (left) and bamboo shark fin buds (right) with scaled TPMs.

There have been several difficulties that limit genetic comparisons between tetrapods and non-tetrapod vertebrates. For example, whereas zebrafish and medaka are ideal models for molecular studies, as they can be genetically engineered, their rapid evolutionary speed and a teleost-specific whole-genome duplication hinder comparative analyses with tetrapods at both the morphological and genetic levels ^11^. This obstacle can be circumvented by using more slowly evolving species such as spotted gar, coelacanths, and elephantfish with their genome sequences that have not experienced recent lineage-specific genome duplications and thus facilitate the tracing of the evolution of gene regulation^12,13^. However, the major disadvantage of these slowly evolving species is the inaccessibility of developing embryos. In contrast, although the eggs of sharks and rays (other slowly evolving species^14^) are often more accessible, their genomic sequence information has not been available until recently. As a solution for these problems, this study used embryos of the brown-banded bamboo shark (referred to hereafter as the bamboo shark), because a usable genome assembly was recently published for this species^14^. Importantly, its non-coding sequences seem to be more comparable with those of tetrapods than with teleosts^14^. In addition, this species is common in aquariums, providing an opportunity to study embryogenesis, and there is a detailed developmental staging table for bamboo shark^15^. These unique circumstances of the bamboo shark enabled a more comprehensive study to identify the genetic differences between fins and limbs.

In this study, to identify genetic differences between fins and limbs, we performed RNA sequencing (RNA-seq) analyses of developing bamboo shark fins and mouse limbs. Along with this transcriptomic comparison, we also generated an accurate orthology map between the bamboo shark and mouse. In addition, we applied an assay for transposase-accessible chromatin with high-throughput and chromatin accessibility analysis (ATAC-seq)^16^ across a time series of mouse limb buds, which generated a high-quality data set showing dynamics of open chromatin regions (OCRs; putative enhancers) during limb development. We also analyzed the evolutionary conservation of sequences in these OCRs to gain insights into the gene regulatory changes during the fin-to-limb transition.

## Results

### Comparative transcriptome analysis

To compare the temporal dynamics of gene expression between bamboo shark fin and mouse limb development, we obtained RNA-seq data from a time series of growing fin and limb buds with three replicates (Fig. 1b; Extended Data Fig. 1 for the details of RNA-seq). We selected limb buds from embryonic day (E)09.5 to E12.5 mice because this is the period during which the major segments of the tetrapod limb—the stylopod, zeugopod, and autopod—become apparent. In particular, the presumptive autopod domain, which is a distinct structure in the tetrapod limb, is visually recognizable from E11.5. For the bamboo shark, we selected developing fin stages from as wide a time period as possible (Fig. 1b; Supplementary Table 1 for details of short-read data). To perform fine-scale molecular-level comparison, we annotated its coding genes using BLASTP against several vertebrates (listed in the Materials and Methods) and our custom algorithm. As a result, 16391 genes from 63898 redundant coding transcripts were annotated as orthologous to known genes of vertebrates, among which 11879 genes were orthologous to mouse genes (Table 1 for details of the transcriptome assembly; Extended Data Figs. 1–3 and Supplementary data 1–6 for gene annotations). The quality of the ortholog assignment assessed by examining *Hox* and *Fgf* genes showed that our custom algorithm is more accurate than other methods (Fig. 1c; see Materials and Methods, Supplementary Table 2 and Extended Data Figs. 1–3 for details).

**Table 1.** Assembly statistics of bamboo shark transcriptome.

| Characteristic | Bamboo shark transcriptome | Bamboo shark gene model |
| --- | --- | --- |
| Total number of sequences | 222015 | 34038 |
| Total sequence length (bp) | 195541367 | 36633751 |
| Average length (bp) | 880 | 1076 |
| Maximum length (bp) | 18451 | 108594 |
| N count | 0 | 10208 |
| L50 | 24765 | 5666 |
| N50 length (bp) | 2075 | 1749 |
| Protein coding | 63898 | 34038 |
| Orthology detected | 42552 | 18783 |
| Unique orthologs | 16391 | 15770 |
| Unique orthologs without gene symbols | 4336 | 3412 |
| Unique orthologs only in elephant sharks | 760 | 508 |
| Sequences with no orthology | 20134 | 14654 |

Using this assembly for the bamboo shark and refseq genes for mice, the transcripts per million (TPM) values were calculated (see Extended Data Fig. 4 for other normalization methods) and scaled by setting the highest TPM in each gene of each species to ‘1’ to capture temporal dynamics rather than absolute transcript amounts. Using this transcriptome data set and gene annotation, we first validated our data by analyzing the expression profiles of *Hoxa* and *Hoxd* genes. As expected, we detected the temporal collinearity of these genes in mouse limb transcriptomes; the expression levels of *Hoxd1* to *Hoxd8* are highest at E09.5 and *Hoxd9* to *Hoxd13* are gradually upregulated later (Fig. 1d). A similar profile was observed for *Hoxa* genes (Fig. 1d). As with mouse limb buds, we found a temporal collinearity in the bamboo shark fin transcriptome (Fig. 1d), suggesting that these transcriptomic data cover comparable developmental stages between the two species at least with respect to *Hox* gene regulation.

Next, to find differences in gene regulation between the two species, we performed a gene-by-gene comparison of expression dynamics with hierarchical clustering (Fig. 2a). To find potential candidate genes that contribute to the different morphologies between fins and limbs, we annotated genes with mouse mutant phenotypes (see Supplementary Table 3 for the full list of genes, expressions, and annotations). The result showed that 8257 genes were significantly expressed in only one of these species (“Fin-specific” and “Limb-specific” in Fig. 2a; 4934 and 3323 genes, respectively). While the fin-specific gene group consisted of many uncharacterized genes, it included ones that are known to control only fish fin development^17,18^, such as *And1* (TRINITY_DN62789_c1_g1_i3 in Suplementary Data 5; ortholog of a gar gene, XP_015216565) and *Fgf24* (TRINITY_DN92536_c7_g1_i2 in Suplementary Data 5; ortholog of a gar gene, XP_015199329). In the limb-specific gene group, several interesting genes were listed that exhibit abnormal phenotype in the mouse limb (e.g., *Bmp2, Krt5, Ihh* and *Megf8*). However, the number of these species-specific genes is probably unreliable and overestimated because these groups also contain genes for which their orthology was not assigned correctly. We also detected that 1114 genes were upregulated during late stages of fin/limb development for both species, including genes that are well known to be expressed later during fin/limb development, such as the autopod-related transcription factors *Hoxd13* and *Hoxa13* and the differentiation markers *Col2a1* and *Mef2c* (“Conserved, late” in Fig. 2a). Intriguingly, 4086 genes exhibited a remarkable heterochronic expression profile; their expression levels were highest during the late stages of mouse limb bud development but decreased during the late stages of fin bud development (“Heterochronic” in Fig. 2a). For validation, we examined the spatio-temporal expression pattern of two heterochronic genes that exhibit limb abnormality in mouse mutants, *Hand2* and *Vcan.* Their transcriptions were upregulated in mouse forelimb buds at E12.5 and downregulated in bamboo shark fin buds at st. 32 (Fig. 2b, c). These results suggest mass heterochronic shifts in gene expression between bamboo shark fin buds and mouse forelimb buds.

**Fig. 2.**
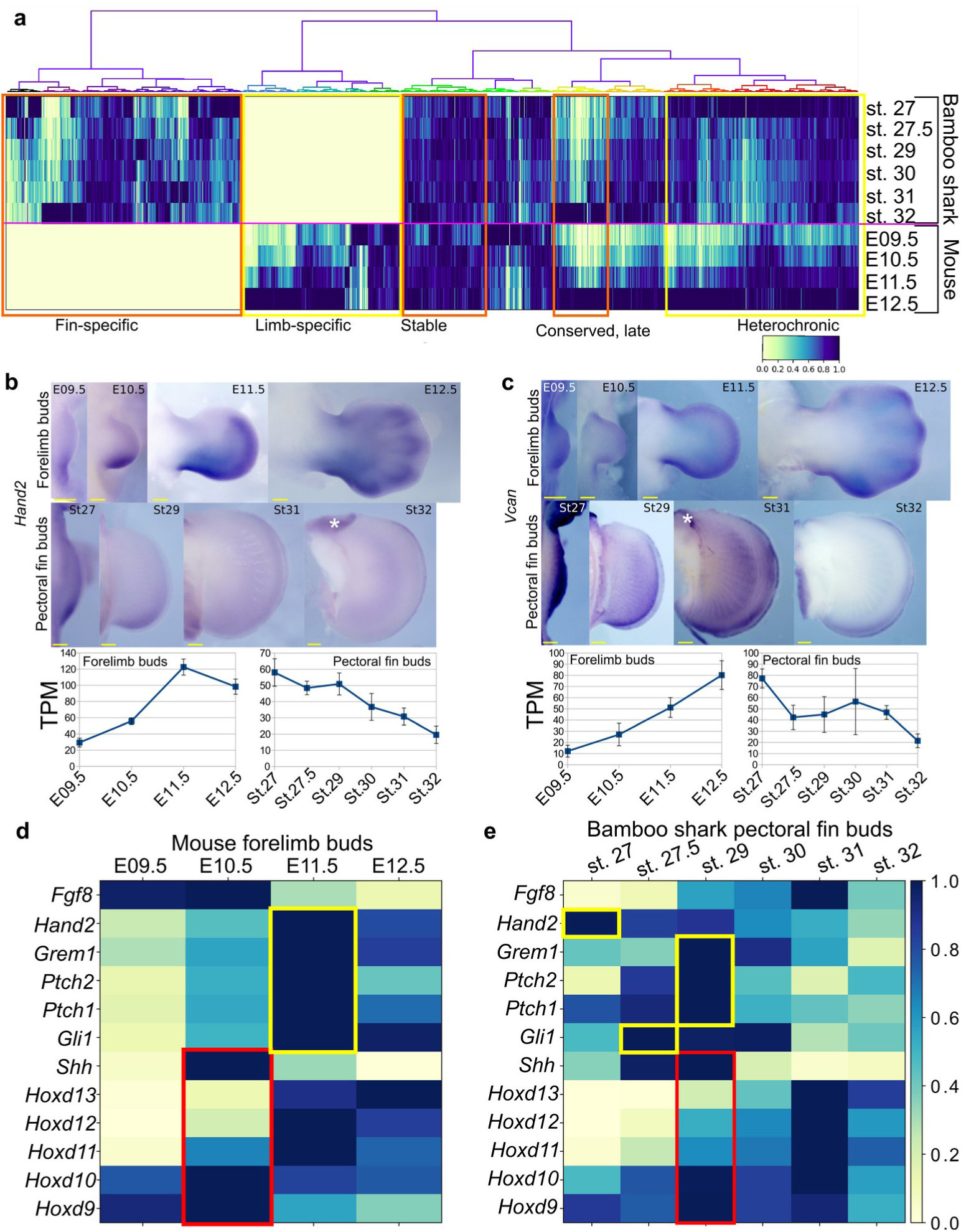
Detection of heterochronic gene expression between mouse limb buds and bamboo shark fin buds. **a**, Clustering analysis of gene expression dynamics. Each column represents an ortholog pair between the bamboo shark and the mouse. Each row indicates scaled gene expression at a time point indicated to the right of the heat map. Values are scaled TPMs. **b, c**, Whole-mount *in situ* hybridization of *Hand2* (**b**) and *Vcan* (**c**) as examples of the heterochronic genes detected in **a**. Asterisks, background signals; scale bars, 200 μm. **d, e**, Expression of *Shh* and related genes in mouse limb buds (**d**) and bamboo shark fin buds (**e**), respectively. The rectangles indicate the expression peaks of *Shh, Hoxd9*, and *Hoxd10* (red) and Shh target genes (yellow).

### Comparison of SHH signaling pathways in limb and fin buds

In tetrapod limbs, SHH controls growth and asymmetric gene expression along the anterior-posterior axis. Previous studies suggested a relatively delayed onset of *Shh* expression or a short signal duration in developing fins of several elasmobranch species^19–21^. Because the heterochronic genes identified above include basic SHH target genes, such as *Ptch1* and *Gli1*, we reexamined the expression dynamics of *Shh* and its target genes in mouse limb and bamboo shark fin buds. Because HOX genes are the upstream factors relative to *Shh* transcription^22^, we used them as a potential reference for developmental time. We first found that *Shh* transcription was present by the earliest stages examined in both bamboo shark fin and mouse limb buds, and it peaked when the transcription level of *Hoxd9* and *Hoxd10* was highest, suggesting that there was no apparent heterochrony in *Shh* transcription timing at least between these two species (red rectangles in Fig. 2d). In contrast, SHH target genes, such as *Ptch1/2, Gli1, Gremlin* and *Hand2*^*23*^, did show a relatively extended period of expression in mouse limb buds as compared with their expression in bamboo shark fin buds. Namely, whereas the expression peak of SHH target genes was concurrent with that of *Shh* in the bamboo shark fin bud, these genes were highly expressed in E11.5 limb buds, which is one day later than the *Shh* expression peak (yellow rectangles in Fig. 2d). We cannot completely reject the possibility that this timing difference is due to the different physical time-resolution of data sampling between these species (six time points over 20 days in the bamboo shark and four time points over 4 days in the mouse). However, given that this data set captured the similar expression dynamics of HoxA/D clusters between these species, the result quite likely represents an interesting difference in the transcriptional regulation of SHH downstream genes between fins and limbs.

### Hourglass-shaped conservation

Several studies have reported a temporally heterogeneous diversification of embryonic transcriptomes, such that the middle stages are more conserved than early or late stages, e.g.,^24–26^. These observations are considered to support the notion of the developmental hourglass (or egg timer), which has been proposed to explain the morphological similarity of mid-stage embryos based on developmental constraints, such as strong interactions between tissues or Hox-dependent organization of the body axis^27,28^. In addition, a previous transcriptomic analysis reported that the late stage of mammalian limb development has experienced relatively rapid evolution^29^. To examine which developmental stages of fins and limbs are conserved, we calculated the distance between the fin and limb transcriptome data. As a result, four different distance methods that we examined consistently indicated that the limb bud at E10.5 and the fin buds at st. 27.5–30 tended to have a relatively similar expression profile (Fig. 3a). In addition, the transcriptomic profile of all the stages of examined fin buds showed the highest similarity to that of E10.5 limb bud (Fig. 3b). Therefore, the mid-stages of limb and fin buds tend to be conserved over 400 million years of evolution.

**Fig. 3.**
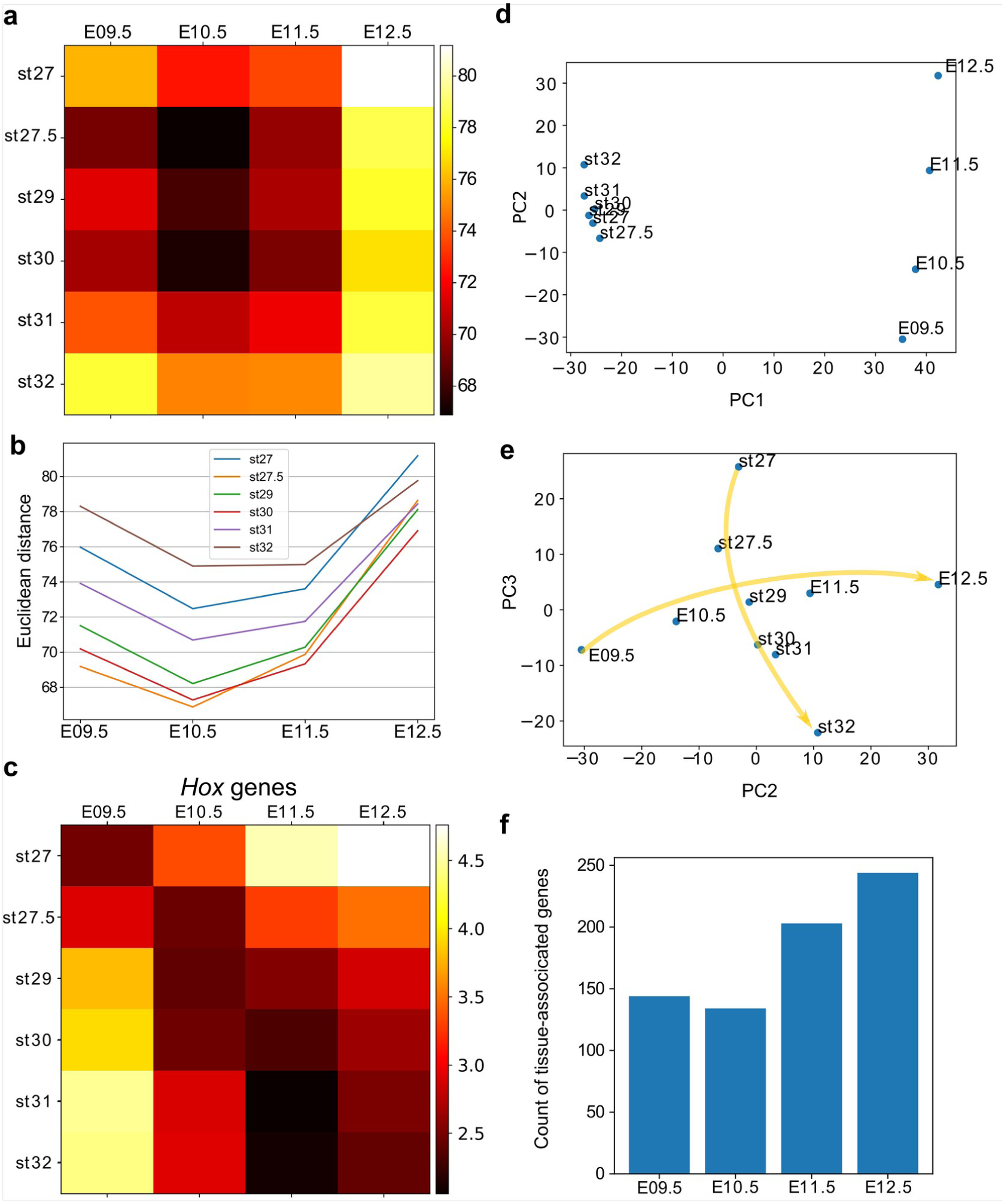
Hourglass-shaped conservation of the transcriptome profile between fins and limbs. **a**, Euclidean distances of the transcriptome profiles. Every combination of time points of bamboo shark fin buds and mouse limb buds is shown. The darker colors indicate a greater similarity between gene expression profiles. **b**, A line plot of the Euclidean distances shown in (**a**). The *x* axis indicates the mouse limb stages, and the *y* axis is the Euclidean distance. **c**, The same as (**a**) except that only *Hoxd* genes are included. **d, e**, Scatter plots of the first and second principal components (**d**) and of the second and third components (**e**). Arrows in (**e**) indicate the time-order of transcriptome data. **f**, Count of tissue-associated genes expressed in mouse forelimb buds. Genes with 0.65 ≤ entropy were counted.

To find factors that underlie the mid-stage conservation, we analyzed *Hox* genes, which were proposed to be responsible for the developmental hourglass^27^. We found that the comparison of only *Hox* gene expression did not reproduce the hourglass-shaped conservation (Fig. 3c), suggesting that other mechanisms constrain the middle stage of development. We further performed principal component analysis (PCA) of gene expression profiles to identify genes responsible for the hourglass-shaped conservation. The first component, PC1, distinguished transcriptome data mostly by species differences (Fig. 3d). In contrast, PC2 was correlated with the temporal order of mouse limb buds (Fig. 3d). PC2 was also weakly correlated with the temporal order of bamboo shark fin buds (Fig. 3d), but PC3 showed a clearer correlation (Fig. 3e). Interestingly, the plot with PC2 and PC3 roughly mirrored the hourglass-shaped conservation because the earliest and latest stages were placed more distantly than the middle stages in this representation (Fig. 3e). The major loadings of these components indicated that the heterochronically regulated genes (e.g., *Tubb4a* and *Tmem200a*) identified in the previous analysis appeared to at least partly contribute to the distant relationship between the early/late stages of fins and limbs (Supplementary data 7). These results indicate that the mass heterochronic shift in gene expression, at least in part, contributes to the long distances between early- and late-stage expression profiles (Fig. 3e).

Because a recent report suggests that pleiotropy of genes is related to hourglass-shaped conservation^30^, we counted the number of genes with stage- or tissue-specific expression. Consistent with the previous report^30^, we detected a relatively low number of stage-associated genes during the middle stages of mouse forelimb and bamboo shark fin development (Extended Data Fig. 5). To evaluate the tissue specificity of genes, we first calculated Shannon entropy of gene expression patterns by analyzing RNA-seq data from 71 mouse tissues as released by the ENCODE project^31^. Namely, genes expressed only in a few tissues score lower with respect to entropy (thus, these genes are more specific). We counted genes with 1.0 ≥ TPM and 0.65 ≤ entropy and, again, found that the number of tissue-associated genes was relatively low at E10.5 (Fig. 3f). Together, these results indicate an inverse correlation between the hourglass-shaped conservation and the number of tissue- and stage-specific genes.

### Open chromatin region (OCR) conservation

Next, we systematically compared gene regulatory sequences between fins and limbs and sought a possible cause for the hourglass-shaped conservation in gene regulatory sequences. To this end, we applied ATAC-seq, which detects OCRs (putative active regulatory sequences), to time-series of forelimb buds at E09.5–E12.5 with three replicates. First, as a positive control, we found that ATAC-seq peaks that were determined by MACS2 peak caller covered 10 of 11 known limb enhancers of the HoxA cluster (Fig. 4a and Extended Data Fig. 6), suggesting a high coverage of true regulatory sequences. Consistently, our ATAC-seq data showed relatively high scores for a quality control index, fraction of reads in peaks (FRiP), as compared with data downloaded from the ENCODE project^31^ (Fig. 4b). Next, to examine evolutionary conservation, we performed BLASTN^32^ for the sequences in the ATAC-seq peaks against several vertebrate genomes. Reinforcing the result of the transcriptome analysis, we found that evolutionarily conserved sequences were most active at E10.5 (Fig. 4c). To confirm this result, we also used a different alignment algorithm, LAST^33^ with the bamboo shark and the alligator^34^ genomes. Alignment results for both analyses consistently indicated that the OCRs of E10.5 forelimb bud more frequently contained conserved sequences relative to those of other time points (Fig. 4d). Therefore, activation of conserved gene regulatory sequences may be one of the proximate causes for the hourglass-shaped conservation of fin and limb transcriptome data.

**Fig. 4.**
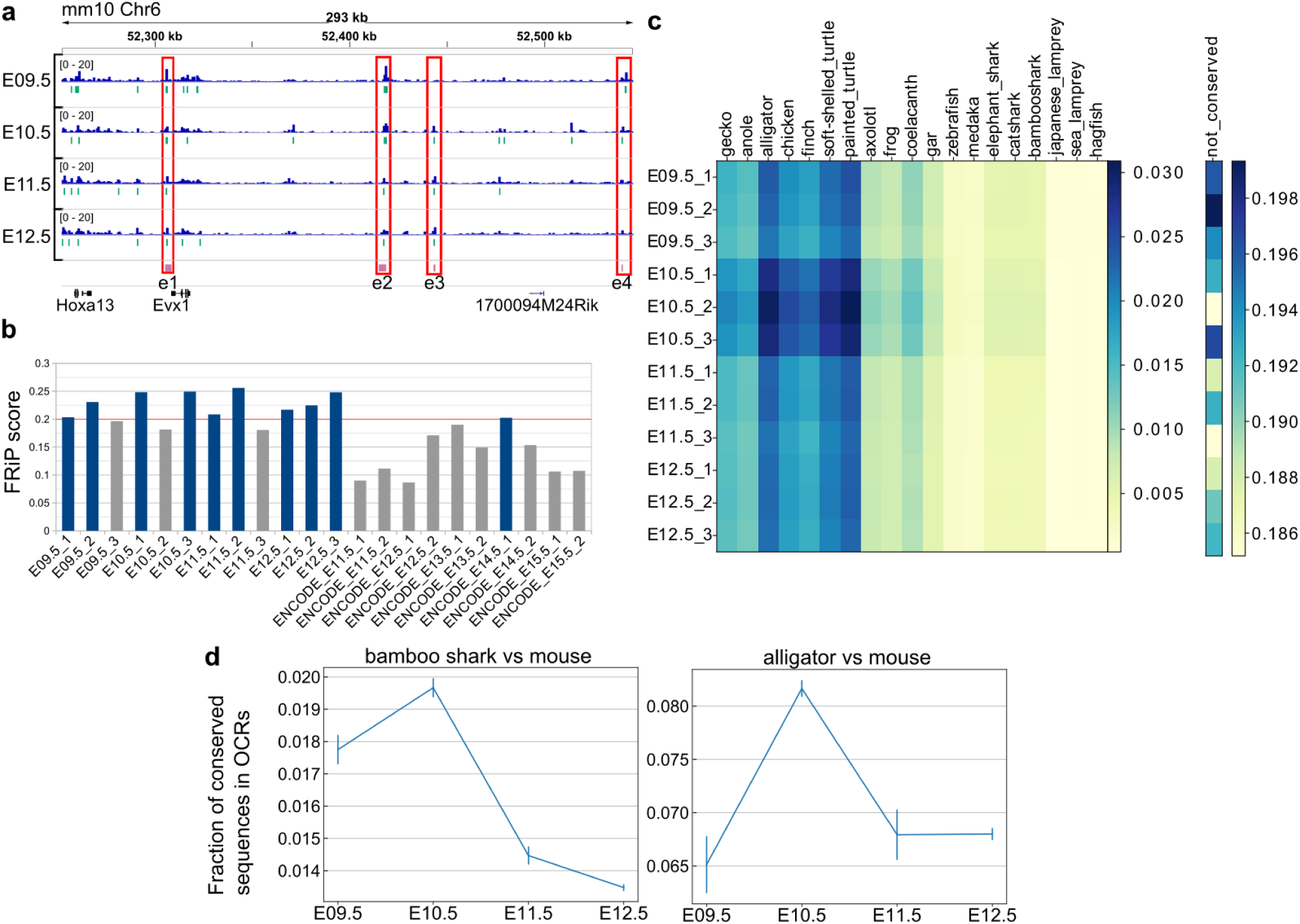
Hourglass-shaped conservation of OCRs in mouse limb development. **a**, ATAC-seq signals in the enhancer regions of the HoxA cluster. e1 to e4, known limb enhancers. Green vertical lines below the signals, peak regions determined by MACS2. **b**, Comparison of a quality index, FRiP, for ATAC-seq data. Blue bars are samples with a FRiP score > 0.2. The number in the end of the label name indicates the replicate number. **c**, Conservation analysis of sequences in ATAC-seq peaks with BLASTP. The values to the right of each graph indicate the fraction of conserved sequences in the total peak regions. The common name of each genome sequence is indicated above the graph. The not-conserved heatmap indicates the fraction of sequences that were not aligned to any genome sequences and thus serve as a negative control. **d**, Temporal changes of sequence conservation frequency in ATAC-seq peaks with LAST.

### Temporal dynamics of open chromatin domains

To further characterize the ATAC-seq peaks, we next performed a clustering analysis. Using one of the three replicates for each stage, we collected the summits of peaks and the surrounding 1400 bp and carried out hierarchical clustering, which resulted in eight clusters (C1–C8; Fig. 5a) that consisted of broad (C1 and C2), E11.5/E12.5-specific (C3 and C4), stable (C5 and C6), E10.5-specific (C7), and E09.5-specific (C8) peaks. The overall clustering pattern was reproducible by other combinations of replicates if its FRiP was ≥ 0.20. Extraction of enriched motifs in each cluster with HOMER^35^ revealed that each cluster contained a specific sequence signature. In particular, it was convincing that stable peaks (C5 and C6) were enriched for motifs for the binding of CTCF, which is a major regulator of three-dimensional genomic structure, and that E11.5/E12.5-specific peaks (C3 and C4) were enriched for HOX13 binding motifs. Interestingly, in E10.5-specific peaks (C7), the LHX1 binding motif was ranked at the top of the motif enrichment list (the closely related transcription factors *Lhx2, Lhx9*, and *Lmx1b* are required to mediate a signaling feedback loop between ectoderm and mesenchyme in limb development^36^). With volcano plots, we also determined the genomic regions that showed a statistically significant increase or decrease in the ATAC-seq signal within a day. As a result, ATAC-seq signals were most increased during the transition from E09.5 to E10.5 in the mouse limb bud. From E10.5 to E11.5, the total number of decreased and increased signals was highest, indicating that the OCR landscape was most dynamically changing at E10.5 (Fig. 5b). In contrast, relatively few significant changes were observed from E11.5 to E12.5. Thus, in contrast to the transcriptome analysis, stage-specific gene regulatory sequences are most active at E10.5.

**Fig. 5.**
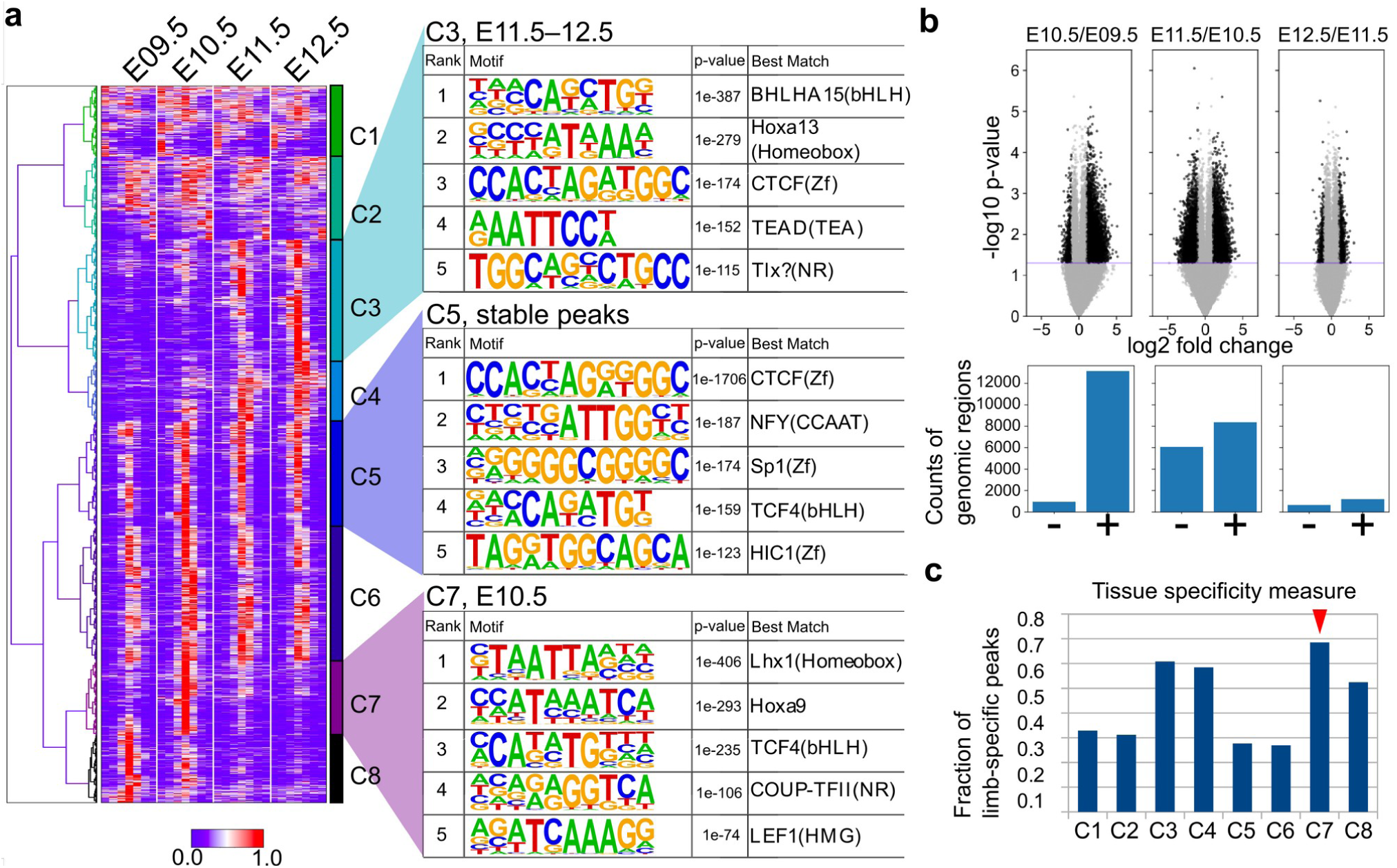
Temporal dynamics of OCRs during limb development. **a**, The heatmap (left) shows whole-genome clustering of ATAC-seq peaks. Each row indicates a particular genome region with a length of 1400 bp. Columns indicate developmental stages. C1–C8 are cluster numbers. The motifs (right) show the rank of enriched motifs in the sequences of each cluster. **b**, Top, volcano plots of ATAC-seq signals between indicated stages (p-values, two-sided Student’s t-test). Bottom, the counts of differential signals (black dots in the top panel). + and − are genomic regions with increased or decreased signals, respectively. **c**, The fraction of limb-specific OCRs for each cluster.

Furthermore, by comparing the peaks of each cluster identified above with ATAC-seq peaks of other cells and tissues released by the ENCODE project^31^ (Supplementary Table 4 for the full list of cells and tissues), we discovered that the C7 cluster (E10.5-specific peaks) contained more peaks that did not overlap with those of other cells and tissues. Again, in contrast to the transcriptome analysis, the data suggest that gene regulatory sequences that are active only at E10.5 tend to be limb-specific (Fig. 5c). Taken together, these analyses revealed a unique regulatory landscape of forelimb buds at E10.5, which is enriched for evolutionarily conserved stage-specific and tissue-specific OCRs.

## Discussions

In this work, we applied transcriptomics and chromatin accessibility analysis to systematically study genetic changes that differentiate fins from limbs. Because of the slow sequence evolution and the embryo availability of the bamboo shark, we were able to compare transcriptional regulation of genes with high accuracy and found both heterochronic shifts and hourglass-shaped conservation of transcriptional regulation between fin and limb development. Here, we discuss the interpretations, limitations, and implications of these results.

Our comparison of time-series transcriptome data indicated that a remarkable number of genes that exhibit weak expression in late-stage fin buds are strongly upregulated in late-stage limb buds. The simplest hypothesis for this mass heterochronic shift is that the later stages of limb development gained expression of one or a few upstream transcription factors or signaling molecules that collectively regulate this group of genes. Interestingly, we also observed relatively extensive expression of the downstream targets of the SHH signaling pathway in mouse limb buds, as compared with bamboo shark fin buds. Because SHH-independent regulation of its target genes through the GLI3-HOX complex was previously reported^37^, the mismatch between the peak expression of *Shh* and its target genes may be caused by such SHH-independent regulatory mechanisms that are absent in bamboo shark fin development. Given that direct and genetic interactions of GLI3 and HOX have a significant impact on autopod formation, the emergence of this interaction may be a key component of the mass heterochronic shift and the acquisition of autopod-related developmental regulation in the tetrapod lineages. However, because we compared only two species, it is equally possible that the late stages of shark fin development lost this gene regulation. Alternatively, given that the evolutionary distance between these two species is >400 million years, it is also possible that every one of these genes independently shifted their expression to the later stages of limb development or to the early stages of shark fin development. Further taxon sampling and functional analyses will reveal the relation between the mass heterochronic shift and the emergence of the autopod.

We observed that gene expression profiles are most highly conserved between bamboo shark fin buds at st. 27.5–30 and mouse forelimb buds at E10.5. Consistent with this result, our chromatin accessibility analysis reveals that OCRs at E10.5 tend to contain evolutionarily conserved sequences. Although transcriptomic and enhancer conservation during the middle of embryonic development have been reported (e.g., ^24,25,38^), as far as we know, this study is the first to convincingly show a correlation between both types of data. Our results suggest that evolutionary constraints on the gene regulatory apparatus are present during the middle stage of fin and limb development. The cause of the hourglass-shaped conservation is still under debate. Interestingly, we found that stage- and tissue-specific OCRs were enriched in this conserved period, during which a relatively low number of stage- and tissue-specific genes were expressed. These quite contrasting observations imply that the mid-stage limb development is enriched for pleiotropic genes controlled by multiple tissue-specific enhancers, including limb-specific ones, rather than by constitutive promoters that often regulate housekeeping genes. Therefore, we speculate that, at least in the case of limb development, complex regulatory sequences that execute spatiotemporally specific transcriptional controls over pleiotropic genes constrain the evolvability of this particular period of morphogenesis, probably due to the vulnerability of complex regulation to genetic mutations.

This study provides a valuable resource not only for comparative studies of fins and limbs but also for the field of limb development and limb-associated diseases, as we provide transcriptome and high-quality open-chromatin data across limb development with a 1-day window. Whereas genome-wide association studies have revealed a number of non-coding mutations associated with human phenotypes^39^, developmental processes are less susceptible to mutations in gene regulatory sequences^40^. This discrepancy suggests that vulnerability to mutations differs among regulatory sequences for unknown reasons. We hope that our detailed catalogue of the open-chromatin landscape of limb development will facilitate understanding of the roles of gene regulatory sequences.

In conclusion, the present work provides insights for the evolutionary origin of gene regulation that differentiates fins from limbs. In particular, comparative transcriptional analyses prompted us to hypothesize that mass heterochronic shifts of gene expression may have occurred during the fin-to-limb evolution. In addition, both transcriptome and open-chromatin data point to an evolutionary constraint during mid-stage limb development, likely owing to gene regulatory complexity. Although these hypotheses require further taxon sampling and experimental tests, this work opens up new prospects for understanding not only the genetic basis of the fin-to-limb transition but also the general nature of morphological evolution.

## Materials and Methods

### Animals

Animal experiments were conducted in accordance with the guidelines approved by the Institutional Animal Care and Use Committee (IACUC), RIKEN Kobe Branch, and experiments involving mice were approved by IACUC (K2017-ER032). The eggs of brown-banded bamboo shark (*C. punctatum*) were kindly provided by Osaka Aquarium Kaiyukan and were incubated at 25°C in artificial seawater (MARINE ART Hi, Tomita Pharmaceutical Co., Ltd.) and staged according to the published staging table^15^. For mouse embryos, C52BL/6 timed-pregnant females were supplied by the animal facility of Kobe RIKEN, LARGE and sacrificed at different days after 9.5−12.5 days of gestation. For RNA-seq, fin buds and limb buds were dissected in cold seawater and phosphate-buffered saline (PBS), respectively, and stored at −80°C. For *in situ* hybridization, embryos were fixed overnight in 4% paraformaldehyde in PBS, dehydrated in a graded methanol series, and stored in 100% methanol at −20°C.

### RNA-seq

Total RNA from mouse forelimb buds and bamboo shark pectoral fin buds was extracted with the RNeasy Micro and Mini plus kit (QIAGEN, Cat. No. 74034 and 74134). Genomic DNA was removed with gDNA Eliminator columns included with this kit. For quality control, the Agilent 2100 Bioanalyzer system and Agilent RNA 6000 Nano Kit (Agilent, Cat. No. 5067-1511) were used to measure the RNA integrity number for each sample. Using 237 ng of each of the RNA samples, strand-specific single-end RNA-seq libraries were prepared with the TruSeq Stranded mRNA LT Sample Prep Kit (Illumina, Cat. No. RS-122-2101 and/or RS-122-2102). For purification, we applied 1.8× (after end repair) and 1.0× (after adapter ligation and PCR) volumes of Agencourt AMPure XP (Beckman Coulter, Cat. No. A63880). The optimal number of PCR cycles for library amplification was determined by a preliminary quantitative PCR using KAPA HiFi HotStart Real-Time Library Amplification Kit (KAPA, Cat. No. KK2702) and was estimated to be 11 cycles for mouse limb buds and 10 cycles for bamboo shark fin buds. The quality of the libraries was checked by Agilent 4200 TapeStation High Sensitivity D1000. The libraries were sequenced after on-board cluster generation for 80 cycles using 1× HiSeq Rapid SBS Kit v2 (Illumina, Cat. No. FC-402-4022) and HiSeq SR Rapid Cluster Kit v2 (Illumina, Cat. No. GD-402-4002) on a HiSeq 1500 (Illumina) operated by HiSeq Control Software v2.2.58 (Run type: SR80 bp). The output was processed with Illumina RTA v1.18.64 for base-calling and with bcl2fastq v1.8.4 for de-multiplexing. Quality control of the obtained fastq files for individual libraries was performed with FASTQC v0.11.5.

### Transcriptome assembly and orthology assignment

We used the NCBI refseq mouse proteins (GRCm38.p5; only curated proteins were used) and two bamboo shark gene lists: a genome sequence–based gene model^14^ and transcripts assembled from RNA-seq in this study (see below) for orthology assignment. The amino acid sequences of the published gene model of the bamboo shark are available from https://figshare.com/articles/brownbanded_bamboo_shark_peptide_sequences_predicted_on_Cpunctatum_v1_0/6125030 (data S1). For the transcriptome assembly, the short reads from the bamboo shark RNA-seq data were trimmed and filtered with Trim Galore! (https://www.bioinformatics.babraham.ac.uk/projects/trim_galore/) and assembled using Trinity^41^ v2.4.0 (options: --SS_lib_type RF --normalize_max_read_cov 200 --min_kmer_cov 2). Protein coding sequences were predicted with a program that finds coding regions, TransDecoder^42^ v3.0.1, according to the guide in TransDecoder (Supplementary data 2 and 3). Using these coding gene lists as queries, orthologous pairs were assigned as illustrated in fig. S1. The idea behind this algorithm is the “gar bridge”^13^, an empirical observation that a comparison including intermediate and slowly evolving animals yields a better resolution for identifying homologous sequences than a direct comparison between two species. First, BLASTP v2.7.1 was performed between mouse and bamboo shark genes reciprocally, and also against the coding genes of the elephant shark (Callorhinchus_milii-6.1.3), spotted gar (LepOcu1), coelacanth (LatCha1), chicken (GRCg6a), alligator (ASM28112v4), and human (GRCh38.p12; options: -outfmt 6 -evalue 1e-30 -window 0). Then, the BLASTP results of bamboo shark queries against the animals listed above (except for the elephant shark) were concatenated, and the best hit across species (cross-species best hit) was identified for each of the bamboo shark genes. If there was no cross-species best hit, then the best hit among the elephant shark genes was retrieved, which may include cartilaginous fish–specific genes. Subsequently, orthologous pairs between mouse and bamboo shark genes were assigned by checking if a cross-species best hit from the bamboo shark BLASTP results also had a best hit in the BLASTP result of mouse genes against the corresponding animal (species-wise best hit; Supplementary data 4–6).

For quality control, the orthology of Fgf family members was independently determined by generating molecular phylogenetic trees (Extended Data Fig. 2 and 3). Amino acid sequences were aligned with an alignment tool, MAFFT^43^ (v7.419-1; options: --localpair --maxiterate 1000) and trimmed with trimAL^44^ (v1.2; options: -gt 0.9 -cons 60). Then, maximum-likelihood trees were constructed with RAxML^45^ (v8.2.12; options: -x 12345 -p 12345 -m PROTGAMMAWAG -f a -# 100). The orthology of *Hox* genes was confirmed based on their synteny. These independently confirmed orthologous pairs were compared with the results of the above orthology assignment algorithm. For a comparison, we also used the results from a reciprocal best hit algorithm, proteinOrtho v6.0.4^46^ and the previously generated orthology groups^14^ (Fig. 1b).

### Quantification and other downstream analyses of transcriptome data

The trimmed RNA-seq short reads were aligned to the transcript contigs for the bamboo shark and curated refseq genes (GRCm38.p5) for the mouse using RSEM v1.3.0^47^ and perl scripts (align_and_estimate_abundance.pl and abundance_estimates_to_matrix.pl) in the Trinity package. TPM (transcripts per million), but not TMM (trimmed mean of M-values), was used for all analyses, because we found some artificial biases in TMM values (see Extended Data Fig. 4). TPM values from the splicing variants of a single gene were summed up to generate a single value per gene. For clustering and distance measures, TPM values were scaled so that the maximum value of each gene of each species was set to ‘1’. Genes with a maximum TPM < 1.0 were considered not expressed. The expression values of each orthologous pair were concatenated as a 10-dimensional vector (consisting of four stages for mouse limb buds and six stages for bamboo shark fin buds), and all gene expression vectors were clustered by t-SNE (hyper parameters: perplexity = 30.000000, n_iter = 5000) followed by hierarchical clustering (hyper parameters: method = “ward”, metric = “euclidean”; the code is available at https://github.com/koonimaru/easy_heatmapper). For the distance measurements, four different distance methods were calculated: Euclidean distance 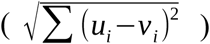, correlation distance 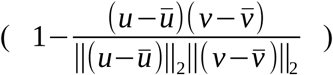, Shannon distance 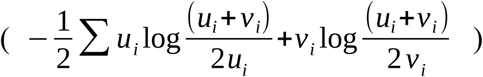, standardized Euclidean distance 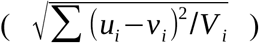, where *u* and *v* are gene expression vectors of two samples and *V*_*i*_ is the variance computed over all the values of gene *i*. For PCA analysis, we used the PCA module in a python package, scikit-learn (https://scikit-learn.org/stable/).

For the stage-associated gene analysis in Extended Data Figs. 5b and 5c, we first calculated the z-score of each gene at each stage as 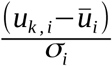, where *u*_*k, i*_ is the TPM value of gene *i* at stage *k, ū*_*i*_ is a mean of TPM over all the stages, and *σ*_*i*_ is the standard deviation of the TPM. Genes with TPM ≥ 1.0 and the absolute Z-score ≥1.0 were counted as stage-associated genes. For the tissue-associated gene analysis, the entropy of each gene was calculated using RNA-seq data of 71 tissues downloaded from the ENCODE web site (https://www.encodeproject.org/; see Supplementary Table 4 for all list). Entropy was calculated as follows:

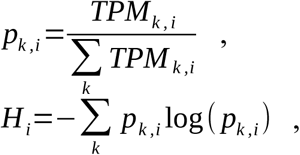

where *TPM*_*k, i*_ is a TPM value of gene *i* in tissue *k, p*_*k, i*_ is a probability distribution and *H*_*i*_ is entropy. Genes with TPM (of mouse limb buds) ≥ 1.0 and 0.65 ≤ entropy were counted as tissue-associated genes.

### WISH

To clone DNA sequences for RNA probes, we used primers that were based on the nucleotide sequences in the ENSEMBL database (https://www.ensembl.org) for mouse genes and in the transcriptome assembly (Supplementary data 3): bamboo shark *Hand2* (Chipun0000087104 in Supplementary data 3), 5′*-*ACCAGCTACATTGCCTACCTCATGGAC-3′ and 5′*-* CACTTGTTGAACGGAAGTGCACAAGTC-3′; bamboo shark *Vcan* (Chipun0000140550 in data S3), 5′*-*AGCTTGGGAAGATGCAGAGAAGGAATG-3′ and 5′*-* AGAGCAGCTTCACAATGCAGTCTCTGG-3′; mouse *Hand2* (ENSMUST00000040104.4), 5′*-* ACCAAACTCTCCAAGATCAAGACACTG-3′ and 5′*-* TTGAATACTTACAATGTTTACACCTTC-3′; mouse *Vcan* (ENSMUST00000109546.8), 5′*-* TGCAAAGATGGTTTCATTCAGCGACAC-3′ and 5′*-* ACACGTGCAGAGACCTGCAAGATGCTG-3′. Fixed embryos were processed for *in situ* hybridization as described^48^ with slight modifications. Briefly, embryos were re-hydrated with 50% MeOH in PBST (0.01% Tween 20 in PBS) and with PBST for 5–30 min each at room temperature (RT). Then, embryos were treated with 20 μg/ml proteinase K (Roche) in PBST (5 sec for mouse E11.5 and E12.5 embryos, 5 min for st. 27 and st. 29 bamboo shark embryos, 10 min for st. 31 and st. 32 bamboo shark embryos). After the proteinase treatment, embryos were fixed in 4% paraformaldehyde/PBS for 1 hour, followed by one or two washes with PBST for 5–10 min each. Optionally, if embryos had some pigmentation, they were immersed in 2% H_2_O_2_ until they became white. Then, embryos were incubated for 1 hour in preheated hybridization buffer (50 ml formaldehyde; 25 ml 20× SSC, pH 5.0; 100 μl 50 mg/ml yeast torula RNA; 100 μl 50 mg/ml heparin; 1 ml 0.5 M EDTA; 2.5 ml 10% Tween 20; 5 g dextran sulfate; and DEPC-treated MilliQ water to a final volume of 100 ml) at 68°C. Subsequently, embryos were incubated with fresh hybridization buffer containing 0.25–4 μl/ml of RNA probes at 68°C overnight. Embryos were washed twice with preheated Wash buffer 1 (50 ml formaldehyde; 25 ml 20× SSC, pH 5.0; 2.5 ml 10% Tween 20; and DEPC-treated MilliQ water to a final volume of 100 ml) for 1 hour each at 68°C; once with preheated Wash buffer 2, which consisted of equal volumes of Wash buffer 1 and 2× SSCT (10 ml 20× SSC, pH 7.0; 1 ml 10% Tween 20; and MilliQ water to a final volume of 100 ml), for 10 min at 68°C; once with preheated 2× SSCT at 68°C for 10 min; and once with TBST at room temperature for 10 min. Embryos were then incubated with a blocking buffer (20 μl/ml 10% bovine serum albumin, 20 μl/ml heat-inactivated fetal bovine serum in TBST) for 1 hour at room temperature, followed by incubation with 1/4000 anti-digoxigenin (Roche) in fresh blocking buffer at 4°C overnight. Embryos were washed four times with TBST for 10–20 min each and were incubated at 4°C overnight. Finally, embryos were incubated with NTMT (200 μl 5 M NaCl; 1 ml 1 M Tris-HCl, pH 9.8; 500 μl 1 M MgCl_2_; 100 μl 10% Tween 20; and MilliQ water to a final volume of 10 ml) for 20 min and then with 15 μg/ml nitro-blue tetrazolium chloride (NBT) and 175 μg/ml 5-bromo-4-chloro-3-indolyphosphate *p*-toluidine salt (BCIP) in NTMT for 10 min to 2 hours until signals appeared. Pictures were taken with an Olympus microscope.

### ATAC-seq

Mouse forelimb buds of each stage were dissected and treated with collagenase for 10 min at room temperature. The tissues were then dissociated into single-cell suspensions by pipetting the mixture and passing it through a 40-μm mesh filter (Funakoshi, Cat. No. HT-AMS-14002); the cell suspension was frozen in CryoStor medium (STEMCELL Technologies, Cat. No. ST07930) with Mr. Frosty (Thermo Scientific, Cat. No. 5100-0001) at −80°C overnight, according to^49^. An ATAC-seq library was prepared as described^16^ with some minor modifications. For library preparation, stored cells were thawed in a 38°C water bath and centrifuged at 500*g* for 5 min at 4°C, which was followed by a wash using 50 μl of cold PBS and a second centrifugation at 500*g* for 5 min at 4°C. Ten thousand cells per sample were collected, without distinguishing dead cells, and were lysed using 50 μl of cold lysis buffer (10 mM Tris-HCl, pH 7.4; 10 mM NaCl; 3 mM MgCl_2_; and 0.1% IGEPAL CA-630). Immediately after lysis, cells were spun at 1000*g* for 10 min at 4°C, and the supernatant was discarded. For the transposition reaction, cells were re-suspended in the transposase reaction mix (25 μl 2× TD buffer, 2.5 μl Tn5 transposase [in the Nextera DNA Sample Preparation Kit, Illumina, Cat. No. FC-121-1031], and 22.5 μl nuclease-free water) and incubated for 30 min at 37°C. The reaction mix was purified using DNA Clean & Concentrator-5 (Zymo Research, Cat. No. D4014) by adding 350 μl of DNA binding buffer and eluting in a volume of 10 μl. After a five-cycle pre-PCR amplification, the optimal number of PCR cycles was determined by a preliminary PCR using KAPA HiFi HotStart Real-Time Library Amplification Kit and was estimated to be four cycles. The PCR products were purified using 1.8× volumes of Agencourt AMPure XP. As a control, 50 ng of mouse genomic DNA was also transposed following the standard procedure of the Nextera DNA Sample Preparation Kit. Sequencing with HiSeq X was outsourced to Macrogen, Inc., which was carried out with HiSeq Control Software 3.3.76 (Run type: PE151bp). The output was processed with Illumina RTA 2.7.6 for base-calling and with bcl2fastq 2.15.0 for de-multiplexing. Quality control of the obtained fastq files for individual libraries was performed with FASTQC v0.11.5.

### ATAC-seq data analysis

The short-read data from ATAC-seq were trimmed and filtered with Trim-Galore! (v0.5.0; options: --paired --phred33 -e 0.1 -q 30). We also removed reads that originated from mitochondrial genome contamination by mapping reads to the mouse mitochondrial genome using bowtie2 v2.3.4.1^50^. The rest of the reads were mapped onto the mouse genome (mm10) using bwa v0.7.17 with the “mem” option^51^. Among the mapped reads, we removed reads with length > 320 bp to reduce noise. The rest of the reads were further down-sampled to around 83.2 million reads to equalize the sequence depth of every sample. Peak calls were done with MACS2 v2.1.1^52^ (options: --nomodel --shift -100 --extsize 200 -f BAMPE -g mm -B -q 0.01; the genomic reads were used as a control for all samples).

For the conservation analysis, the significant variation in the length of ATAC-seq peaks complicated this evaluation. To deal with such variation, we first divided the mouse genome into 100-bp bins. Then, the ATAC-seq peaks were re-distributed into these bins with bedtools^53^ (options: intersect -F 0.4 -f 0.4 -e -wo). Peaks of >100 bp were subdivided into 100-bp-long regions, and those of <100 bp were extended to fit within the closest 100 bp window. The sequences in these peaks were retrieved with BLASTN v2.7.1 against 19 genomes of vertebrate species listed in table S4 (BLASTN options: -task dc-megablast -max_target_seqs 1). The blast hits that scored >40 were considered as conserved sequences. In this way, the final figures shown in Fig. 4C represent the fraction of the total conserved sequence length in the peaks of each stage rather than the number of conserved peaks. For confirmation, we also used a different alignment algorithm, LAST v961^33^ to find conserved sequences. To generate mouse genome databases for LAST, we first masked repeat sequences with N and split the genome file into multiple files, each of which contained a single chromosome sequence. Then, databases were generated using lastdb (options: -cR01). Alignments with the bamboo shark genome (Cpunctatum_v1.0; https://transcriptome.riken.jp/squalomix/resources/01.GCA_003427335.1_Cpunctatum_v1.0_genomic.rn.fna.gz) and the alligator genome (ASM28112v3) were carried out by lastal (options: -a1 - m100). Only a unique best alignment was selected using last-split. These alignment results were converted into the bed format, and regions that overlapped with the ATAC-seq peaks that were subdivided into 100-bp bins were counted.

For the clustering analysis, we converted the alignment files of the ATAC-seq reads into mapped reads in bins per million (BPM) coverage values with 200-bp resolution using bamCoverage in deepTools^54^ v3.2.1 (options: -of bedgraph --normalizeUsing BPM -- effectiveGenomeSize 2652783500 -e -bs 200). Then, BPMs at the summits of ATAC-seq peaks and an additional 600 bp to the left and to the right of each summit (1400 bp in total) were collected and clustered by t-SNE (hyper parameters: perplexity = 30.0, n_iter = 5000) followed by hierarchical clustering (hyper parameters: method = “ward”, metric = “euclidean”). Enriched motifs were detected using a perl script, findMotifsGenome.pl in HOMER v4.10.4^35^ (http://homer.ucsd.edu/homer/motif/). For the tissue-specificity analysis, we downloaded several aligned and unaligned reads from the ENCODE web site (https://www.encodeproject.org/; see Supplementary Table 4 for a complete list), and peaks were called as described above. Then, peaks that did not overlap with other tissues/cells were detected using bedtools.

## Acknowledgments

We thank Kenta Sumiyama and James Shape for fruitful discussions; Itoshi Nikaido and Laboratory for Bioinformatics Research for providing computational resources; Itsuki Kiyatake, Kiyonori Nishida, and Osaka Aquarium Kaiyukan for kindly providing bamboo shark eggs; the animal facility of Kobe RIKEN, LARGE for supplying mouse embryos; the ENCODE consortium and the ENCODE production laboratories for generating ATAC-seq datasets.

## Author contributions

KO and SK designed the project. KO, KT, CT, and MK performed experiments. KO and ON analyzed data. All authors contributed to writing the manuscript.

## Data and materials availability

RNA-seq and ATAC-seq datasets generated during the current study are available in the Gene Expression Omnibus (GEO) repository under accession number GSE136445. Code for clustering analysis is available at https://github.com/koonimaru/easy_heatmapper. Additional data related to this paper may be requested from the authors.

## Supplementary Materials

**Extended Data Figure 1 to 6 (below)**

**Supplementary Table 1 to 4\*\***

**Supplementary data 1 to 7\*\*\***

**\*\***found in separate files that accompany this manuscript.

**\*\*\***found in https://figshare.com/articles/Onimaru_et_al_Supplementary_Data/9928541.

****Other NGS-related data are available at GSE136445 (https://www.ncbi.nlm.nih.gov/geo/query/acc.cgi?acc=GSE136445).

**Extended Data Fig. 1.**
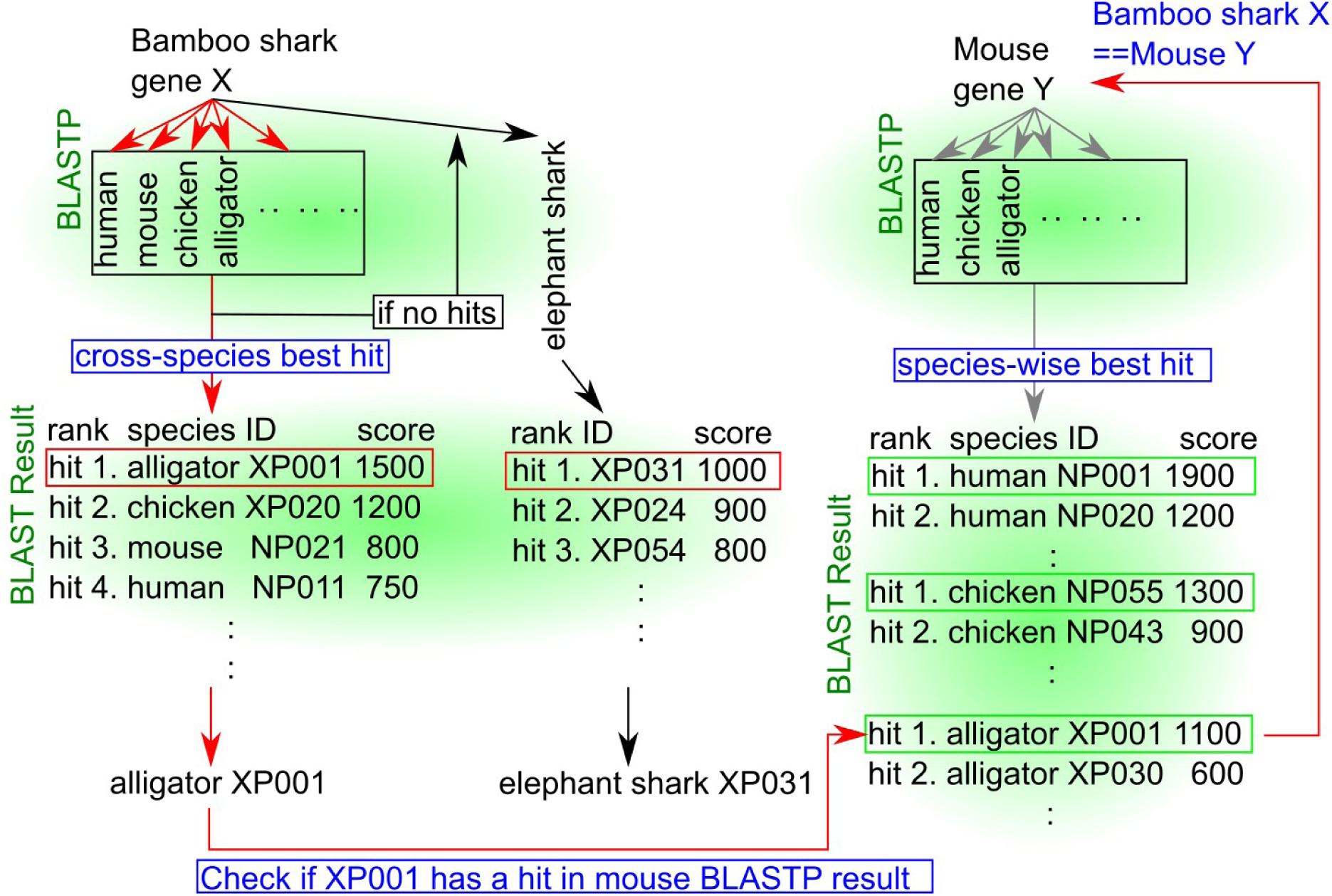
Schematic representation of the orthology assignment algorithm. Red arrows, the main flow of the algorithm. Black arrows, orthology assignment for cartilaginous fish-specific genes. Gray arrows, parallel retrieving of orthologs of mouse genes from other animals. Red rectangles, best hits across other animal genes or in elephant shark genes. Green rectangles, best hits among each animal genome. Note that this schematic explains how the orthology of abstract genes “bamboo shark gene X” and “mouse gene Y” are assigned. First, using BLASTP, putative orthologs of bamboo shark genes are retrieved from other animal genomes, such as human, mouse, alligator, and elephant shark. Then, all BLASTP results except those from elephant shark are concatenated to find a best scored gene across species (cross-species best hit). In this schematic, the alligator gene XP001 is the best hit. In parallel, putative orthologs of mouse genes are also retrieved from the same set of animal genomes. If there is a mouse gene Y that has a best hit with alligator XP001, this mouse gene Y and bamboo shark gene X are considered to be an orthologous pair.

**Extended Data Fig. 2.**
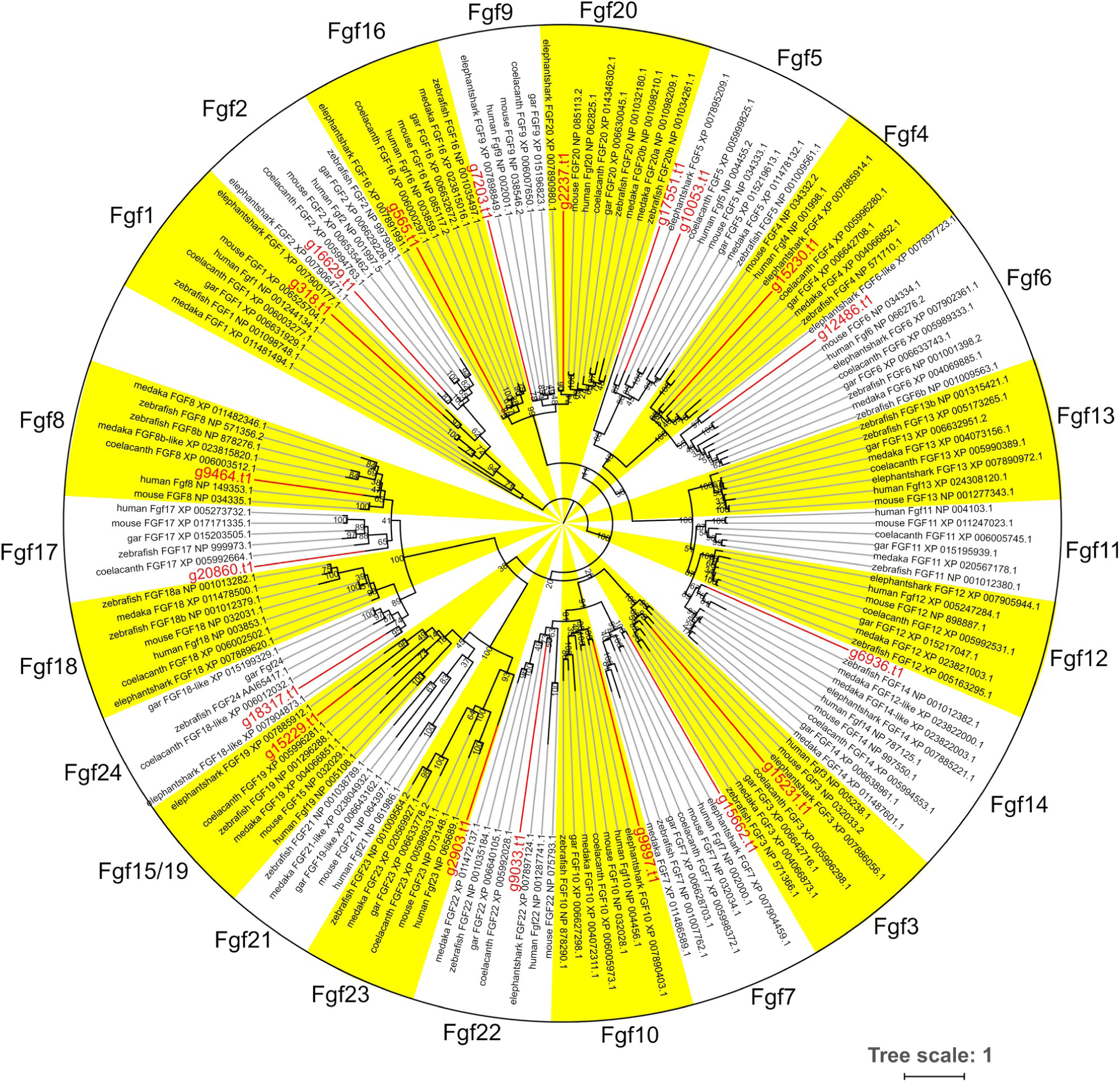
Molecular phylogenetic tree for Fgf family. The tree was inferred with the maximum-likelihood method. The support values at nodes indicate bootstrap probabilities. Genes highlighted in red are bamboo shark genes (can be converted into the original gene ID by replacing “g” with “Chipu000”).

**Extended Data Fig. 3.**
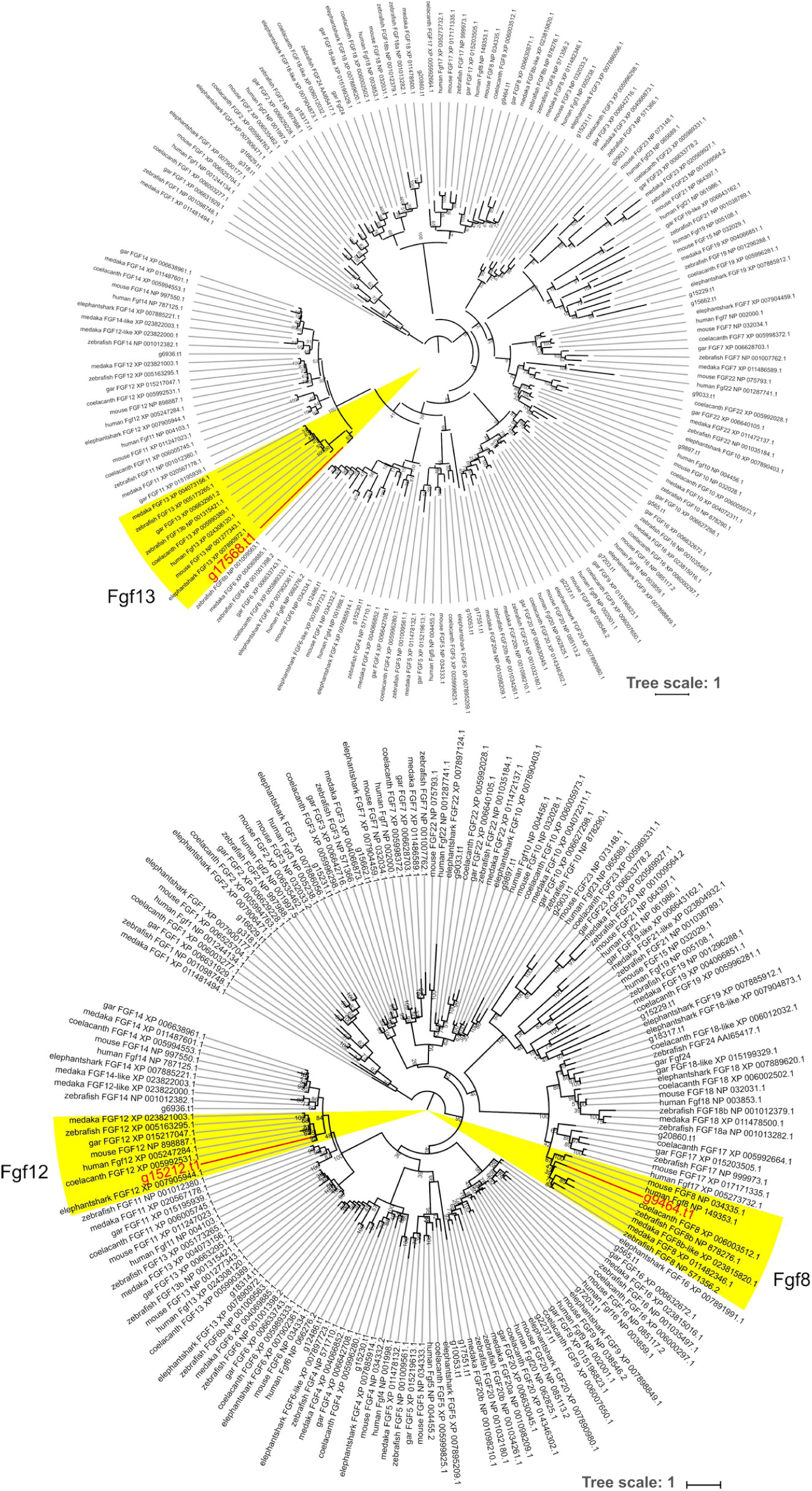
Additional molecular phylogenetic trees for Fgf8, Fgf12, and Fgf13. These trees are included because alignment sequences used in Fig. S2 are truncated or absent in these genes. The tree was inferred with the maximum-likelihood method. The support values at nodes indicate bootstrap probabilities. Genes highlighted in red are bamboo shark genes.

**Extended Data Fig. 4.**
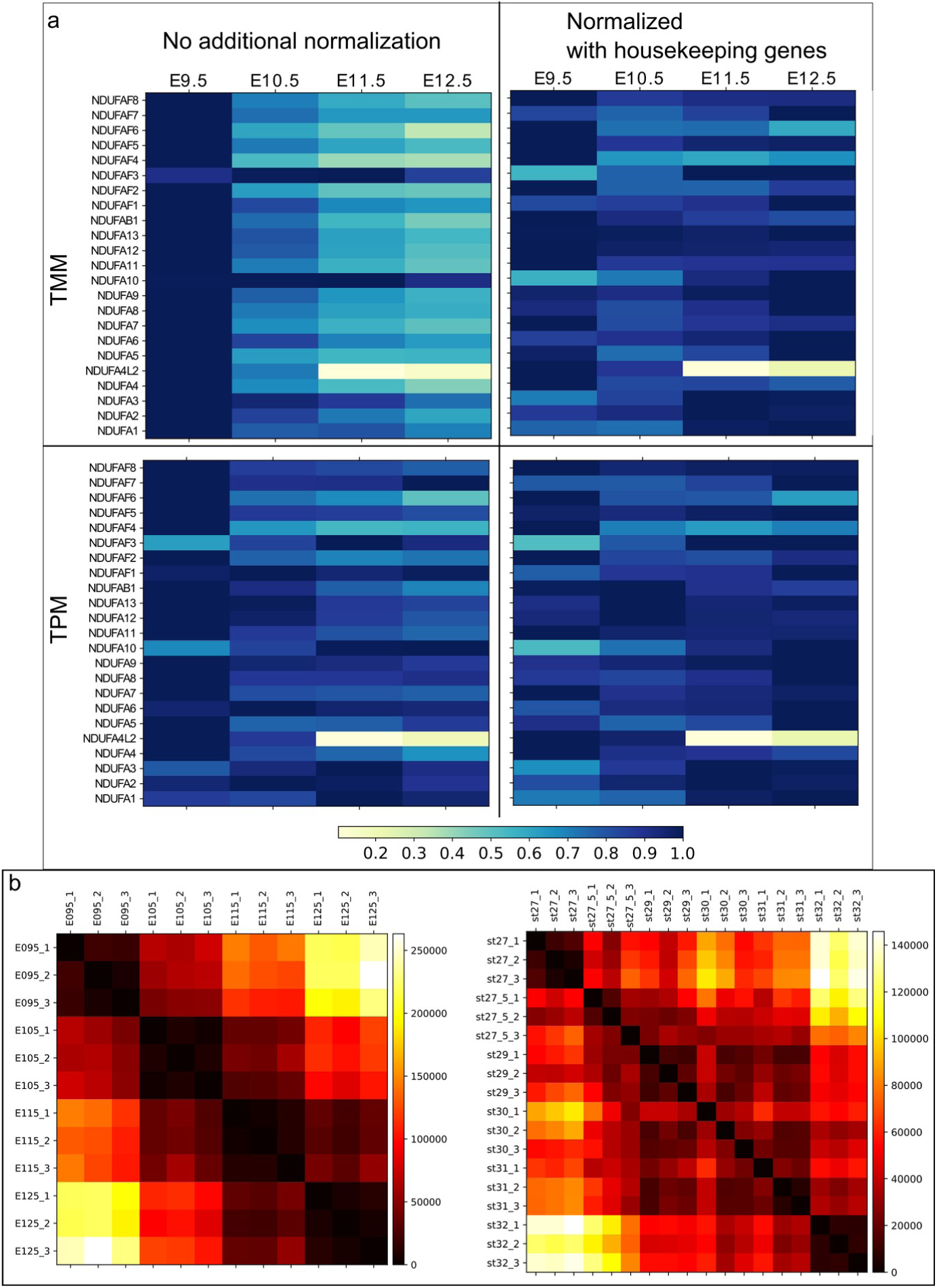
Supplementary data for RNA-seq analysis. **a**, Visualization of the effect of normalization by showing a housekeeping gene family, *Ndufa*. Left panels show TMM (trimmed mean of M) and TPM (transcripts per million) calculated by RSEM. Right panels show these values with additional normalization using several other housekeeping genes (*Atp5j, Atp5h, Atp5g3, Psmc3, Psmc5, Psmd7, Mrpl54, Mrpl46, Polr2e, Polr1b, Mrpl2*). All expression values are standardized by setting the maximum expression value of each gene as ‘1’. Note that because housekeeping genes do not change their expression amount over time, these expression values should be close to ‘1’ (i.e., all colors should be dark blue) with some exceptions. However, the intact TMM (top, left panel) is apparently biased, in that the majority of *Ndufa* genes show their strongest expression at E09.5, with sharp decreases at other stages. This bias can be corrected by normalization with other housekeeping genes (top right panel). In contrast, the intact TPM (bottom, left panel) has a weaker bias than TMM. Additional normalization (bottom, right panel) has less of an effect. Therefore, this study used the intact TPM. **b**, Euclidean distances of transcriptome data between mouse samples (left) and between bamboo shark samples (right). Whereas the close relation of the replicates of mouse samples can be seen from this heat map, the replicates of bamboo shark samples show less similarity. This noisy data may be attributed to the fact that there is no established strain of the bamboo shark and/or that bamboo shark embryos were staged by morphology but not physical time. However, the average of replicates seems to mitigate the noise of the bamboo shark samples, because *Hox* gene expression showed a smooth temporal collinearity as seen in Fig. 1d.

**Extended Data Fig. 5.**
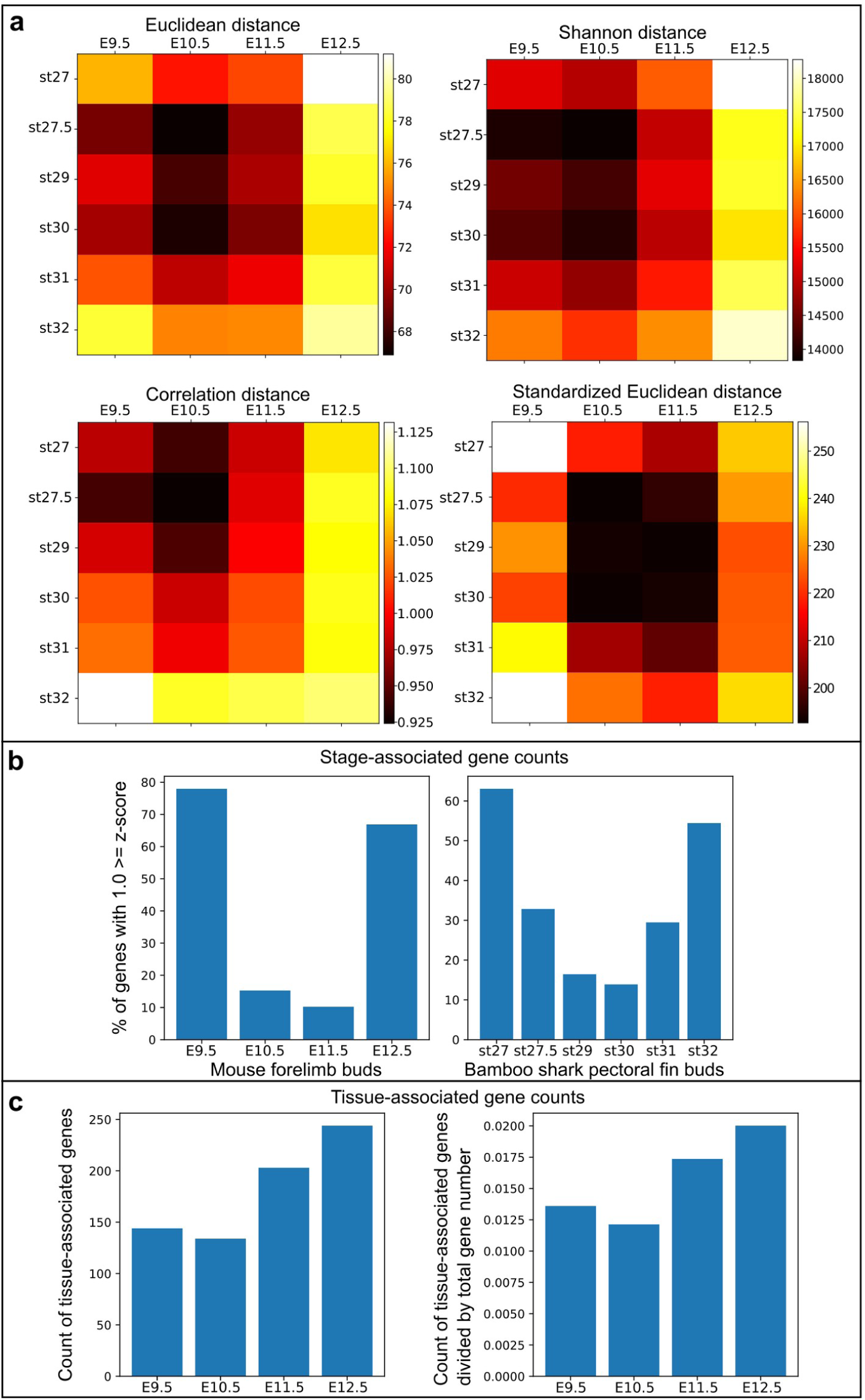
Confirmation analyses of the transcriptome comparison. **a**, Cross-species comparisons of transcriptome data between the two species with indicated distance methods. Note that these methods consistently show the closest distance around E10.5 and st. 27.5– 30. **b**, Percentages of stage-associated genes with 1.0 ≥ z-score of mouse limb buds (left) and bamboo shark fin buds (right). Note that both species show a low percentage of stage-associated genes during the middle stages. **c**, Counts (left) and fractions (right) of tissue-associated genes expressed in mouse limb buds. Tissue specificity was evaluated by entropy using RNA-seq data from 71 mouse tissues. A gene with 0.65 ≤ entropy was considered a tissue-specific gene. In the right panel, gene counts were normalized based on the counts of total expressed genes. Note that the number of tissue-associated genes was lowest at E10.5.

**Extended Data Fig. 6.**
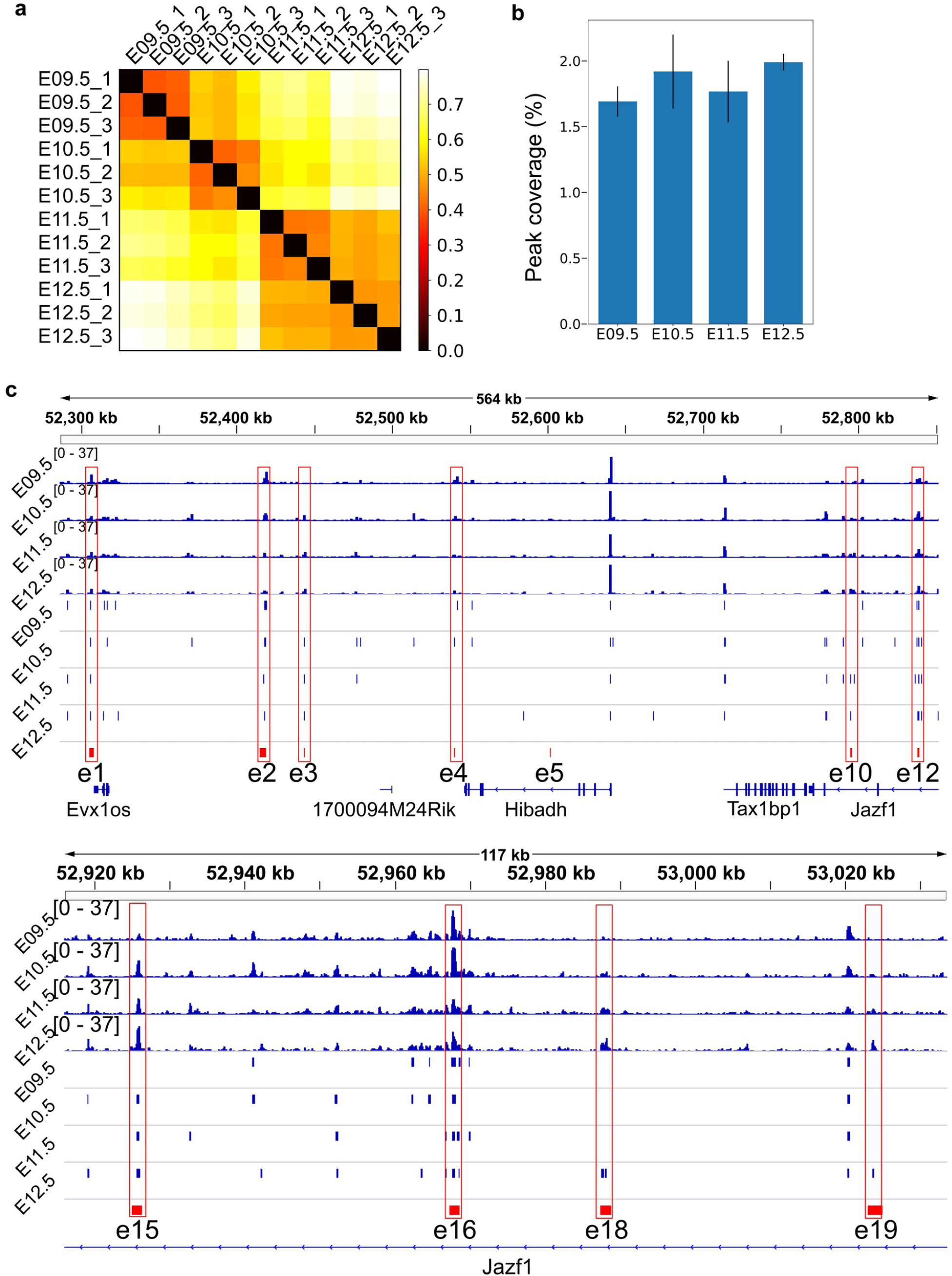
ATAC-seq quality control. **a**, Correlation distance between samples. The numbers in the end of the sample names indicate the replicates of indicated stages. Darker color means more similar gene expressions. **b**, Percentage of peak regions in the genome sequence. **c**, ATAC-seq signals in BPM (blue signals), peak regions (blue rectangles) and the known limb enhancers of HoxA cluster (red rectangles, e1–e19). Note that only e5 is not covered by ATAC-seq data.

